# Potential for Zika virus to establish a sylvatic transmission cycle in the Americas

**DOI:** 10.1101/047175

**Authors:** Benjamin M. Althouse, Nikos Vasilakis, Amadou A. Sall, Mawlouth Diallo, Scott C. Weaver, Kathryn A. Hanley

## Abstract

Zika virus (ZIKV) originated and continues to circulate in a sylvatic transmission cycle between non-human primate hosts and arboreal mosquitoes in tropical Africa. Recently ZIKV invaded the Americas, where it poses a threat to human health, especially to pregnant women and their infants. Here we examine the risk that ZIKV will establish a sylvatic cycle in the Americas, focusing on Brazil. We review the natural history of sylvatic ZIKV and present a mathematical dynamic transmission model to assess the probability of establishment of a sylvatic ZIKV transmission cycle in non-human primates and/or other mammals and arboreal mosquito vectors in Brazil. Brazil is home to multiple species of primates and mosquitoes potentially capable of ZIKV transmission, though direct assessment of host competence (ability to mount viremia sufficient to infect a feeding mosquito) and vector competence (ability to become infected with ZIKV and disseminate and transmit upon subsequent feedings) of New World species is lacking. Modeling reveals a high probability of establishment of sylvatic ZIKV across a large range of biologically plausible parameters. Probability of establishment is dependent on host population sizes and birthrates and ZIKV force of infection, but a network of as few as 6,000 primates with 10,000 mosquitoes is capable of supporting establishment of a ZIKV sylvatic cycle. Research on the susceptibility of New World monkeys or other small mammals to ZIKV, on the vector competence of New World *Aedes, Sabethes*, and *Haemagogus* mosquitoes for ZIKV, and on the geographic range of these species is urgently needed. A sylvatic cycle of ZIKV would make future elimination efforts in the Americas practically impossible, and paints a dire situation for the epidemiology of ZIKV and ending the ongoing outbreak of congenital Zika syndrome.

## Introduction

The invasion of Brazil by Zika virus (ZIKV) is the latest upheaval in a decade-long cataclysm in the epidemiology of viruses transmitted by the mosquito *Aedes aegypti* in the Americas [1, 2]. Dengue virus (DENV) made forays into Florida in 2009 [3], Arizona in 2014 and Hawaii in 2015 [4]; chikungunya virus (CHIKV) was introduced into the Caribbean in 2013 and hurtled across both Central and South America [5]; and, in 2015, Zika virus (ZIKV) was first detected in Brazil. With ZIKV came a spike in cases of congenital microcephaly and Guillain-Barre syndrome [2, 6]. The introduction of ZIKV to the Americas had been predicted well in advance of the event [7, 8], however, the association of ZIKV infection with neuropathology and teratogenicity were only revealed during the spread of the virus through the Pacific and into Brazil. Hayes [7] did warn in 2009 that the spread of ZIKV warranted concern despite lack of contemporary evidence for severe ZIKV disease. He reminded the scientific community that West Nile virus was also considered a “relatively innocuous pathogen” until it ushered outbreaks of neuroinvasive disease into Europe and the Americas. In response to a growing body of evidence linking ZIKV infection with teratogenic effects [9, 10, 11], the World Health Organization declared the ZIKV outbreak a public health emergency of international concern in February of 2016 [1, 12].

ZIKV is unusual among the arthropod-borne viruses (arboviruses) in its capacity for sustained transmission in a human-endemic cycle. This capacity is shared by three other arboviruses that are also, not coincidentally, transmitted in the human cycle by *Aedes aegypti*: DENV, CHIKV and yellow fever virus (YFV). For all four viruses, human-endemic lineages emerged from ecologically and evolutionarily distinct, sylvatic, enzootic cycles transmitted between mostly arboreal *Aedes* spp. vectors and non-human animal hosts [7, 13, 14]. While non-human primates (hereafter primates) have generally been considered the major reservoir hosts for the sylvatic transmission cycle of all four viruses, this paradigm is based on scant evidence and researchers in the field have repeatedly cautioned that other animal species may play key roles in the transmission dynamics of these viruses [15, 16, 13, 7]. The ancestral sylvatic cycles of YFV, CHIKV and ZIKV occur in Africa, while the DENV ancestral cycle occurs in Southeast Asia with later transport to West Africa and enzootic establishment [17]. YFV was transported from Africa to the Americas in infected humans and mosquitoes via the slave trade in the 17th and 18th centuries [18] and spilled back into a sylvatic cycle, maintained in New World monkey species, which persists today. Dengue virus, in contrast, has not spawned an established a sylvatic transmission cycle in the Americas despite widespread circulation of the virus across the Americas in the human-endemic cycle [13].

Whether ZIKV will emulate YFV or DENV is an open and urgent question. If the virus establishes a sylvatic cycle in the Americas, then mosquito control and even vaccination will not suffice to eradicate it from the region. ZIKV was first isolated in 1947 from a sentinel monkey in the Ziika forest of Uganda. Intriguingly the sentinel species used was the rhesus macaque, demonstrating the susceptibility of Asian primates to ZIKV. The next year ZIKV was isolated from *Ae. africanus* in the area, suggesting mosquito-borne transmission of the virus. As laid out in the comprehensive review by Hayes, the virus was subsequently detected across a wide swath of tropical Africa via serosurveys of monkeys as well as virus isolation from monkeys and several species of sylvatic *Aedes* [7]. Notably these mosquitoes were collected in the forest canopy but also on the forest floor. Infection of humans living in proximity to sylvatic cycles was detected via serosurveys and clinical surveillance. Seroprevalence was variable and quite high (up to 40%) in some human populations, but disease was invariably mild, generally manifesting as fever, headache, rash and conjunctivitis. In 2007, an outbreak of ZIKV in Libreville, the capital of Gabon was thought to have been vectored by the peri-urban mosquito species *Aedes albopictus* [19]. Importantly, experimental studies of the interaction among different African arboviruses have shown evidence for both enhancement [20, 21] and interference [7].

Over the same time period that the ZIKV transmission cycle was being investigated in Africa, circulation of ZIKV was documented in several countries in Asia. Albert Rudnick, the pioneer of sylvatic DENV research, isolated the virus from *Ae. aegypti* in Malaysia [22]. A serological study in 1977-78 in Central Java revealed that a high percentage of febrile patients had antibodies against ZIKV [23]. Subsequently ZIKV infection was documented in travellers returning from Indonesia [24], Thailand [25], and Malaysia [26] and in residents of Indonesia [27], Cambodia [28], the Philippines [29], and Thailand [30]. One of the cases of Zika infection in a traveller was notable as disease onset occurred five days after being bitten by a monkey in Indonesia [31]. Anti-ZIKV antibodies have also been detected semi-captive orangutans in Malaysia [32] To date there has been no solid evidence of an Asian sylvatic cycle of ZIKV, but such a sylvatic cycle could be widespread and still go undetected due to the lack of surveillance for sylvatic arboviruses in Southeast Asia [33]. Thus it remains uncertain whether all human ZIKV infections in Asia derive from the human-endemic cycle or whether some may occur due to spillover from an as-yet unknown sylvatic cycle in the region. The lineage of ZIKV that circulates in Asia is distinct from the African lineages of the virus, and it is the Asian lineage that spread across the Pacific and into Brazil [34].

Research on the sylvatic cycle of ZIKV since 2007 has focused primarily on West Africa. Phylogenetic analysis indicates that the virus has been introduced into West Africa at least twice in the twentieth century [35] and that West Africa contains ZIKV strains that are distinct from those elsewhere in Africa [36]. Analyses of mosquitoes collected annually over the last fifty years in Kedougou, Senegal demonstrated that ZIKV is amplified in mosquito collections at four year intervals, that rainfall is a positive predictor of ZIKV isolations in mosquitoes, and that there was little positive or negative association between amplification of ZIKV and of three other *Aedes*-borne arboviruses that circulate in the region, YFV, DENV-2 and CHIKV [37]. Moreover our field studies in Kedougou during the 2011 ZIKV amplification showed that the virus was present in all major land cover classes in the region but was detected significantly more often in the forest than in other land cover types [38]. In this study, ZIKV was detected in ten separate species of *Aedes*, with *Ae. hirsutus, Ae. unilineatus, Ae. metallicus*, and *Ae. africanus* having the highest minimum infection rates of collected species. In addition, one pool of male *Ae. furcifer* was found to be positive, indicating possible vertical transmission of ZIKV. To follow up these field observations, Diagne et al. tested the vector competence of multiple Senegalese *Aedes* species for ZIKV in the laboratory and found that only *Ae. luteocephalus* and *Ae. vittatus* were capable of transmitting the virus [39]. ZIKV has previously been isolated from two of the three monkeys species resident in Kedougou: African green monkeys (*Chlorocebus sabaeus*) and patas monkeys (*Erythrocebus patas*) (reviewed in [40]) In combination with previous field studies in Africa, these findings demonstrate that the transmission dynamics of ZIKV are complex and that a diverse network of *Aedes* vector species and primate host species participate in the maintenance of the sylvatic ZIKV cycle.

Here, we used a mathematical model that we have previously employed to study the sylvatic DENV cycle in Senegal [41] to identify the conditions of host and vector density and connectivity that would permit the establishment of an American sylvatic cycle of ZIKV.

## Establishing a sylvatic ZIKV cycle

Our model extends, to our knowledge, the only previous dynamic model of mosquito-borne viruses in non-human primate hosts [41, 42]. While the Althouse et al. (2012) study was focused on sylvatic DENV, the strong similarities between sylvatic DENV and sylvatic ZIKV transmission cycle make the model a good fit for both viruses. Here we use the model to ask whether ZIKV will establish a self-sustaining transmission cycle in a network of two susceptible host populations with two corresponding competent vector mosquito populations after introduction of a single ZIKV-infected host. We assume host and vector species interact as separate populations, and thus populations correspond to separate species. Further, we assume that each host population has a vector population that is source of the largest number of bites that could transmit ZIKV, indicating vector preference, with a vector biting its preferred host 100 times more frequently than its non-preferred hosts. Our previous work suggests that this non-preferred biting synchronizes transmission in the two populations and the synchrony is qualitatively unrelated to the ratio of preferred to non-preferred biting [41]. Here we explore primates and mosquitoes as the hosts and vectors, with Althouse et al. (2012) and 1Althouse and Hanley (2015) giving full model details.

Briefly, mosquitoes and primates are born susceptible to ZIKV infection, and are infected at a rate proportional to the number of bites given or received per day and a probability of infection. Primate species differ in their life history, particularly birthrate and lifespan. We assume birthrate = 1/lifespan, which is conservative as age of fertility completion is younger than age of mortality for many primates [43]. Transmission probabilities vary seasonally due to differences in rainfall and temperature [37]. We explore three per-bite infection probabilities (0.3, 0.6, 0.9) with an average of 0.5 infectious bites per day. This gives forces of infection 0.15, 0.3, 0.45, which is in line with observed sylvatic DENV forces of infection from primate collections in Kedougou, Senegal in 2010-2012 (Sall, Diallo, Althouse, Hanley, Weaver, unpublished data). These forces of infection ranged from 0.09 (95% CI: 0.07, 0.11) for Guinea baboons (*Papio papio*) in 2012, to 0.41 (95% CI: 0.26, 0.76) for African green monkeys (*Chlorocebus sabaeus*) in 2012. After infection, primates recover at a fixed rate (4 days [44]) while mosquitoes are infected for the remainder of their life. We employ the stochastic version of the model simulated using a Gillespie stochastic simulation algorithm with the Binomial Tau leap approximation (BTL) to examine the effects of population size, primate birthrate, and force of infection on the probability of ZIKV establishment. Simulations were run and we calculated the proportion of simulations not becoming extinct after introduction of a ZIKV infected host (ie, establishing a sylvatic cycle).

Model simulations suggest the probability of establishment is highly dependent on the primate birthrate (Figure 1). In low and medium force of infection settings (0.15 and 0.3) primates with lifespans of 15 and 25 years show little probability of sylvatic establishment (panels d, g, h). However, if there exists a rapidly reproducing primate (lifespan of 5 years), establishment of a sylvatic cycle is nearly assured (panels a, b, c). Generally, increasing numbers of primates relative to mosquitoes lowers the probability of establishment, as might be expected as the force of infection is directly proportional to the number of mosquitoes and inversely proportional to the number of primates [37]. A network of as few as 6,000 primates with 10,000 mosquitoes is capable of supporting the establishment of a ZIKV sylvatic cycle.

**Figure 1:**
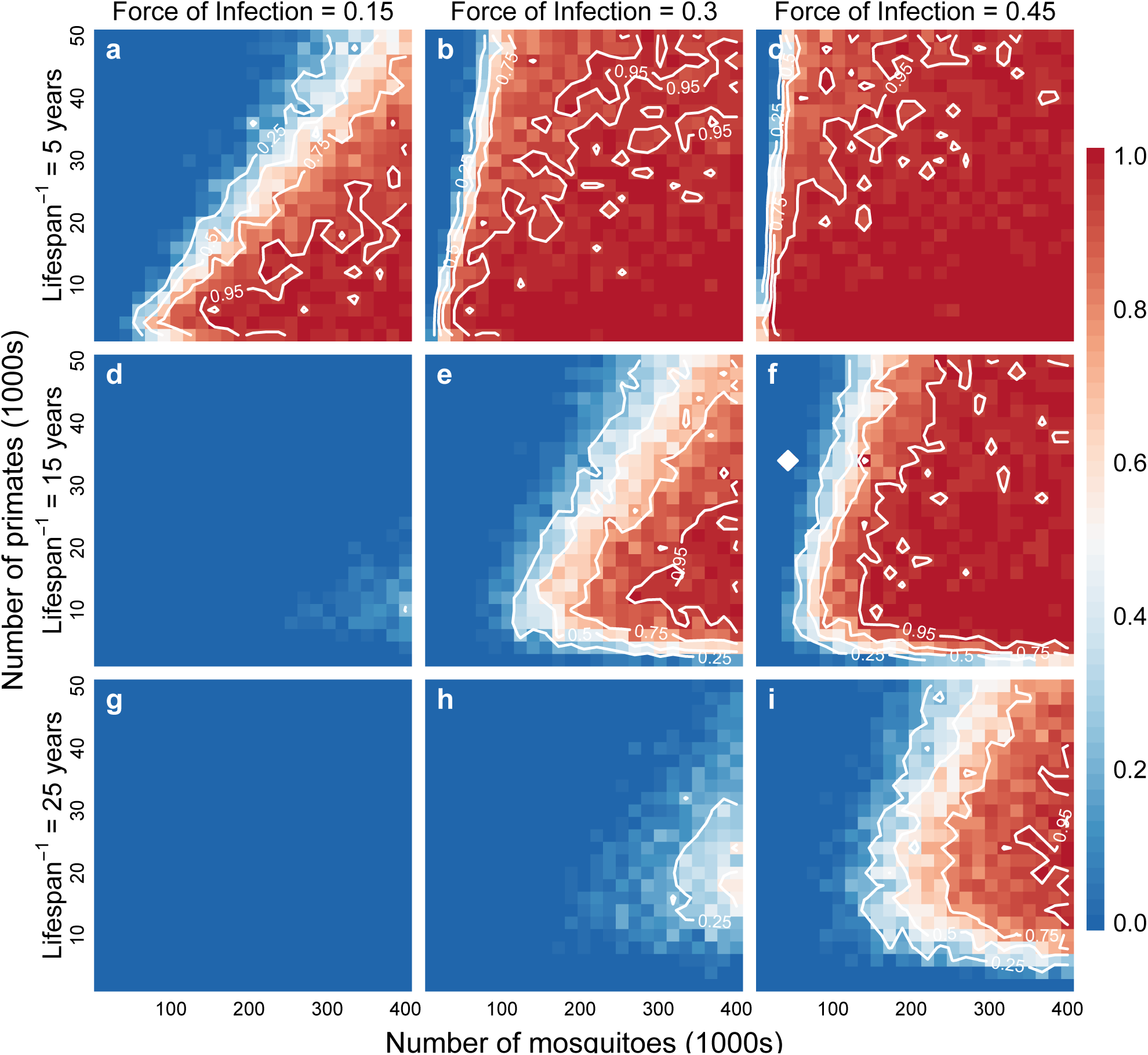
Figure 1 Probability of establishing a sylvatic ZIKV transmission cycle. Figure shows heat maps of the probability of ZIKV establishment in 50 simulations per parameter set with colors ranging from blue (0 simulations establishing) to red (all simulations establishing). Contour lines show 0.25, 0.5, 0.75, and 0.95 probability of establishment. For each plot, the x-axis shows the total number of mosquitoes (in two populations) and the y-axis shows the total number of non-human primates (in two populations). Left to right the panels indicate increasing in force of infection, and top to bottom decreasing non-human primate birthrate (as 1/lifespan^15^). Other parameters: mean mosquito lifespan = 7 days; mean ZIKV recovery in NHP = 4 days; mosquito vertical transmission of ZIKV = 0; rate of yearly ZIKV introduction = 0.

## Outlook

To our knowledge, the susceptibility of New World monkeys to ZIKV has never been tested, and it is possible that they are insusceptible to ZIKV infection or generate only low levels of viremia insufficient to infect potential sylvatic vectors. However, as we have pointed out in a previous review, there are free-living populations of several Old World monkey species in the Americas, some of which, notably African green monkeys (which as noted above had high forces of sylvatic DENV infection in Senegal) are known to be hosts of sylvatic ZIKV in Africa [13]. Our model predicts that the presence of a rapidly reproducing primate or other mammal that is a competent host for ZIKV vastly increases the chances of establishment. There is some serological evidence that vertebrates other than primates may also serve as enzootic reservoirs of ZIKV [27, 45]. Again, ZIKV susceptibility testing of potential small mammal hosts is lacking and should be a high priority for future research.

We also do not know the susceptibility of most New World *Aedes* species for ZIKV. However, it has recently been shown that *Ae. albopictus* was likely the primary vector of a ZIKV outbreak in humans in Gabon [19]. This mosquito species, which is common in the Americas, has a broad host range and has high potential to serve as a bridge vector to transfer the virus from humans to non-human animals [46]. Additionally, *Sabethes* and *Haemagogus* spp mosquitoes are tropical New World vectors of sylvatic YFV and thus may be likely vectors of sylvatic ZIKV as well [13].

The current work is limited by gaps in knowledge, and relies on sylvatic ZIKV transmission dynamics being similar to sylvatic DENV transmission – a reasonable assumption given the extensive overlap of the two viruses in the hosts and vectors used in West African sylvatic cycles [7, 13, 38]. We note that our model calculates the probability of ZIKV establishment starting from a single infectious introduction without further importation, and does not include vertical transmission of ZIKV within mosquitoes. These features both make our estimates conservative and potentially paint a dire situation for the epidemiology of ZIKV and for prospects of extinguishing the ongoing congenital Zika syndrome outbreak in Brazil.

The International Task Force for Disease Eradication identifies a key factor for considering a disease eradicable as epidemiologic vulnerability, including not having the presence of an animal reservoir [47]. Establishment of a sylvatic cycle of ZIKV would make future elimination efforts in the Americas extremely difficult if not impossible. Taking lessons from sylvatic YFV in Brazil, reactive, and massive vaccination efforts will be necessary when a ZIKV vaccine becomes available to control ZIKV transmission [48], decrease morbidity, and protect unborn infants from potential teratogenic effects. We use this work to identify and highlight key lines of research that would enable the public health community to understand ZIKV transmission going forward and target surveillance for enzootic ZIKV to those animal populations most likely to sustain virus transmission.

## Acknowledgements

We greatly acknowledge NSF-RCN on Infectious Disease Evolution Across Scales program. This work was supported by grants 1R15AI113628-01 (KAH), RO1AI069145 (BMA, SCW, KAH), R24AI120942 (SCW) from the U.S. National Institutes of Health. The funders had no role in study design, data collection and analysis, decision to publish, or preparation of the manuscript.

